# The impact of serial translocations on the genetic diversity of Anegada iguanas (*Cyclura pinguis*) in the British Virgin Islands

**DOI:** 10.64898/2026.02.18.705091

**Authors:** Giuliano Colosimo, Zachary Dykema, Mark E. Welch, Gabriele Gentile, Gad Perry, Zachary Harlow, Glenn P. Gerber

## Abstract

Animal translocations are becoming increasingly popular as a tool for conservationists. Demographic factors can be crucial determinants dictating translocation viability in the short term. Translocated populations pass through artificial bottlenecks and can suffer from founder effects. Reduction in genetic variation relative to their source populations is likely, limiting their adaptive potential. Founder events can increase frequencies of deleterious alleles due to elevated rates of inbreeding and inbreeding depression. Here, we describe the effects of human-driven, serial population translocations on the genetic diversity of critically endangered Anegada iguanas (*Cyclura pinguis*) in the British Virgin Islands. Though founding populations were extremely small (N=8, N=4), the census sizes of translocated iguana populations increased dramatically over the first twenty years. This implies that these translocations were successful from a demographic perspective despite the small number of animals used, indicating a genetic paradox. To quantify genetic signatures in these bottlenecked populations, blood samples were collected from the source population and two translocated populations and genotyped at 21 microsatellite loci. We found that allele frequencies in translocated populations differed significantly from those of the source, with the translocated populations having less genetic diversity. However, common methods for estimating presence of genetic bottlenecks were non-significant. Estimates of internal relatedness by age class suggest that inbreeding depression may be elevated after translocation, likely reflecting the small initial population sizes associated with these translocation events. Anecdotally, our work shows that translocations may result in subtle genetic erosion that has long-term population viability impacts, even when census size indicates success.

## Introduction

Conservation translocation is the human-driven movement of organisms from one location to another for the benefit of that species’ survival and of the ecosystem it naturally inhabits (IUCN/SSC 2013). Conservation translocations may involve adding individuals to an existing population (supplementation or reinforcement), moving individuals back to their indigenous range after extirpation (reintroduction), and moving individuals outside of their native range for the benefit of the species as a whole (assisted colonization) or to fill an extinct ecological function (ecological replacement) (IUCN/SSC 2013). Besides supplementation, which essentially acts as assisted migration, all other translocation methods establish a new population of individuals.

Animal translocation has become an increasingly useful tool for the management and conservation of wildlife imperiled by extinction, though results have varied (Berger-Tal et al. 2019; Morris et al. 2021; Evans et al. 2023). The founding of a population in a new habitat is usually smaller than the source population, thus leading to different selection pressures, fitness outcomes, and allele frequency changes from genetic drift (Santos et al. 2012). For a translocation to be considered successful in the long-term, the new populations must be both viable and adaptable. Viability requires that the founders survive, reproduce, and their offspring successfully recruit to the breeding population (Forsman 2014; Szűcs et al. 2014). Long-term success may not be achieved if the nascent population is established with too few individuals, thereby increasing extinction risk. For example, stochasticity with demographic rates, disease, and climactic events have larger impacts on recent colonization due to small population size (Rajakaruna et al. 2015). Ecological mismatch between the source environment and the new environment also may prevent successful colonization due to different abiotic and biotic factors (Pintar & Resetarits 2021, Cardador et al. 2022).

Moving too few organisms can have detrimental genetic effects on the nascent population because it passes through an artificial bottleneck, a reduction in genetic variability as a consequence of reduced population size following translocation (Nei et al. 1975; Templeton 1980; Allendorf 1986; Santos et al. 2012; Hartl 2020). Although bottlenecks are a central focus for conservation biologists, they are not necessarily viewed with the same concern by invasion biologists. Species naturally colonizing new areas and expanding their original habitat via propagule pressure are often able to overcome the hindrances of new populations founded by only a handful of individuals (Kekkonen and Brommer, 2015). This apparent contradiction between conservation and invasion biology gave rise to the so-called “genetic paradox” of invasive and colonizing species (i.e., how newly founded populations overcome low genetic diversity and evolutionary potential to persist in the colonized environment; Allendorf & Lundquist, 2003). Currently this paradox is considered resolved or, rather, difficult to prove. To truly be considered a genetic paradox, in fact, the newly founded population must *i)* have lower genetic variation than the source population, *ii)* not be demographically impacted by said lower genetic variation, and *iii)* adapt well to the new environment it has been placed in (Estoup et al., 2016). However, many supposed genetic paradoxes have comparable or increased genetic variation compared to the native population from multiple introduction events and admixture among founders (Uller & Leimu 2011, Estoup et al., 2016). Moreover, even in the case of documented genetic impoverishment, newly established populations often do not show any significant adaptive challenge and can reproduce successfully if the environment is benign or similar to that of the source population (Estoup et al., 2016). Colonization might also be facilitated by preadaptation or exaptation, the co-option of characteristics that contribute to fitness in a novel environment (Hufbauer et al., 2012).

Regardless of the actual occurrence of a genetic paradox, conservation biologists should prudently monitor the genetics of a population involved in an assisted migration since the dispersal capacities of the species target of a translocation are, by definition, insufficient to colonize new environments, and the translocated population cannot naturally rely on further propagules to augment its genetic variability (Kekkonen and Brommer, 2015). Endangered species, operating as small populations, are limited in their genetic variation (Frankham 2005). The genetic hindrances of a translocation can further be aggravated if multiple translocations are seeded using individuals from a previous translocation event. This chain of multiple bottlenecks can amplify the loss of genetic diversity and may result in population decline, the insurgence of physical abnormalities, and an increase in extinction probability (Gautschi et al. 2002; Leberg and Firmin 2008).

In the present study we explore genetic variation in three populations of Anegada iguanas (*Cyclura pinguis*) to assess the genetic impact of serial bottlenecks resulting from conservation translocations in the British Virgin Islands (BVI). This system is especially valuable since there is a record of source population genetic diversity, when and how many founders were translocated, and no known immigration between islands prior to, or after, translocation. We first reconstruct the history of two translocation events that occurred between 1984 and 1995 (see Materials and Methods for details). We then use microsatellite data to compare genetic variation in the natural and translocated populations. Current estimates of population census size of these translocations indicate successful establishment from a demographic perspective. This could indicate a genetic paradox where the iguana populations have overcome founder effects and preserved their evolution potential. Given the low number of individuals used to seed the two translocation events, however, we predicted that there would be significant differences in allele frequencies between the three populations, with the source population having the greatest genetic diversity. We also evaluated common genetic bottleneck statistical tests and approaches for estimating effective population size to see how these approaches compare to the species’ known demographic history. As more translocations are used to prevent local extinctions and initiate recolonization of species in their natural habitat, we use this case as an example to provide some indication of how this practice may affect the survival of translocated populations.

## Materials and Methods

### Study system

Anegada iguanas, *Cyclura pinguis*, are one of 10 recognized species of rock iguanas (genus *Cyclura*) (Iguana Taxonomy Working Group–ITWG 2016). This species, endemic to the BVI, is listed as Critically Endangered (CR) under the International Union for the Conservation of Nature (IUCN) Red List of Threatened Species™ (Bradley and Grant 2020). It represents the most basal and unique rock iguana lineage, which originated 10–20 million years ago (Reynolds et al. 2022). Thus, the conservation of Anegada iguanas could help maintain more genetic diversity than the conservation of any other single rock iguana species. *C. pinguis* is sexually dimorphic with males larger than females and individuals usually reach sexual maturity at 6 to 7 years and about 29 cm Snout-Vent-Length (SVL) for males and 36 cm SVL for females (Bradley and Grant 2020).

Historic evidence suggests that the range of *C. pinguis* once extended across the Greater Puerto Rican Bank and numbered over 4000 animals (Carey 1975). Major population declines are thought to have occurred from climatic fluctuations, leaving the island of Anegada as the sole habitat refugium for *C. pinguis* (Perry and Gerber 2006; Figure 1). Further declines began in the late 1960s when development on Anegada led to the proliferation of introduced species and desertification. Goats, sheep, donkeys, and cattle compete with iguanas for food resources alongside feral cats, rats and dogs that have a direct impact on the survivorship of young age classes (Goodyear and Lazell 1994; Mitchell 1999). Extant iguanas on Anegada were reduced to approximately 160 individuals by the early 1990s (Mitchell 1999).

**Fig. 1.**
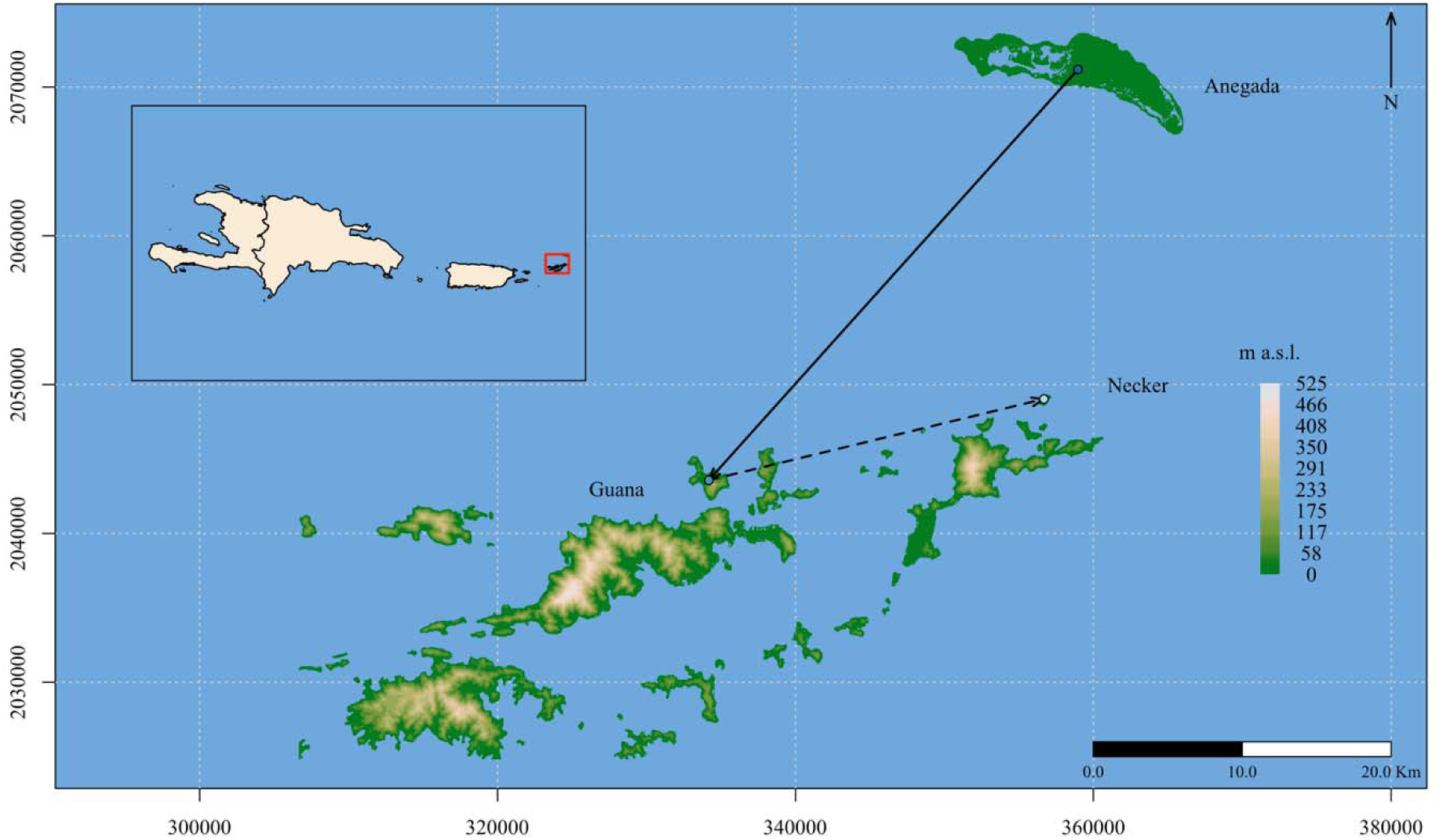
Topographic map of the British Virgin Islands (BVI) with elevation expressed as meters above sea level (m a.s.l.). The relative location of the BVI in the Greater Antilles is highlighted with a red square in the inset map. The main map shows the first (solid line) and second (dashed line) order translocations between Anegada, Guana and Necker islands. Between 1984 and 1986, eight individuals were moved from Anegada to Guana. In 1995, four hatchlings were translocated from Guana to Necker Island. The Guana and Necker populations of iguanas both currently number in the hundreds

Conservation efforts to prevent further declines in the species include three principal strategies. The first is directed at controlling the feral cat population and other invasive mammals on the island. The second focuses on increasing recruitment on Anegada through a headstart and release program that raises juvenile iguanas away from cats until they are large enough to better avoid predation (Perry and Gerber 2006). The third implements translocations by moving animals from Anegada to other islands. The mammal eradication efforts and headstart program are not within the scope of this study, but the slow progress in controlling exotic mammals is being temporarily held in check by successes in the headstart program. However, headstarting juveniles is a temporary solution to the problems faced by this species, and the long-term viability of the species is still very precarious (Bradley and Gerber 2006). Therefore, strategies implying population restoration may serve as a critical tool in managing this species (IUCN/SSC 2013).

Prior to the current conservation actions under consideration, translocations of *C. pinguis* took place. The first translocation occurred between 1984 and 1986 and consisted of eight individuals brought from Anegada to Guana Island (Figure 1). This translocation consisted of three males and five females (224-509 mm SVL), two of which were palpably gravid at the time of release (Goodyear and Lazell 1994). The translocation led to a rapid population increase on the island with an estimated 100 individuals by 2002 (Perry and Mitchell 2002). More recently, this population is estimated to be 300 animals or more (Gerber, pers. obs.). In 1995, nine years after the first translocation, four hatchlings (two females, two males) were moved from Guana Island to Necker Island (Figure 1) and cage reared in captivity until their release the following year (Lazell 2002). The same author reported the survival of all four animals and the emergence of new hatchlings by 1999. As on Guana, the population on Necker has increased expeditiously and currently is in the hundreds (Gerber and Colosimo, pers. obs.). Within the last several decades, translocations of *C. pinguis* to other small private islands have occurred, but little information exists regarding the circumstances of these translocations or their success (Perry and Gerber 2006).

### Data collection and genotyping

A total of 269 individuals were originally considered for this study. Upon capture by either noose or net, each individual was sampled for morphological features, including snout-vent length (SVL), tail length (TL), and mass, as well as sex and age class. We considered hatchlings to have < 100 mm SVL while adults have >250 mm SVL. A passive integrated transponder (PIT, Trovan™, United Kingdom) was applied subcutaneously for long term identification. Additionally, ca. 0.5 mL of blood was collected from the caudal vein using heparinized 1 mL syringes. Blood was stored in EDTA buffer (Longmire et al. 1997) at environmental temperature while in the field prior to long term storage in −80℃ freezers.

Samples were scaled down to 174 as we excluded known hatchling clutch mates in an effort to reduce biases while estimating the distribution of genetic variation. Of the samples considered for this study, 73 individuals were sampled on Anegada, 63 on Guana Island, and 38 on Necker Island (see Table 1 for details). The Anegada specimens were collected between 1999 and 2005. Samples from Guana and Necker were collected between 2006 and 2015. Genomic DNA was extracted from whole blood with Qiagen QIAamp DNA Mini Kit and accompanying handbook protocols. A total of 21 loci were amplified using primers previously developed for *C. pinguis* (Lau et al. 2009, Supplementary Table 1) and following PCR conditions as indicated by Lau and colleagues (2009). Fragment length was determined using an ABI 3100 or 3130xl Genetic Analyzer™ and GENEMAPPER™ Software (Applied Biosystems).

**Table 1.** Sampled Island (Population), number of individuals collected on each site (N), average number of different alleles per locus (N_A_), rarefied allelic richness (A_R_), rarefied private allelic richness (Private_R_), observed heterozygosity (H_o_), and expected heterozygosity (H_e_). These statistics were calculated using GenAlEx 6.5 (Peakall and Smouse, 2012).

| Population | N | $N_A$ | $A_R$ | $Private_R$ | $H_o$ | $H_e$ |
| --- | --- | --- | --- | --- | --- | --- |
| Anegada | 73 | 4.095 | 3.871 | 7.808 | 0.550 | 0.564 |
| Guana | 63 | 3.286 | 3.129 | 0.603 | 0.482 | 0.503 |
| Necker | 38 | 3.143 | 3.056 | 1.000 | 0.454 | 0.444 |

### Data analysis

#### Genetic diversity

We first investigated the presence of scoring mistakes and/or null alleles in our dataset. The misinterpretation of electropherogram peaks during microsatellite genotyping and the presence of null alleles or large allele dropouts can inflate the frequency of homozygous genotypes and bias estimates of overall population diversity and differentiation. We used the software Micro-Checker 2.2.3 (van Oosterhout et al. 2004), which not only evaluates the frequency of null alleles and short allele dominance, but also helps in the detection of scoring mistakes due to microsatellite stuttering and/or other genotyping errors (van Oosterhout et al. 2004). We used all methods for calculating null allele estimates in the software (Chakraborty et al. 1992, Brookfield 1996) We then used the software FreeNA (Chapuis and Estoup 2007; Chapuis et al. 2008), which calculates the frequency of null alleles using the Expectation Maximization (EM) algorithm described in Dempster et al. (1977). We repeated this approach separately for samples from each island.

Unless otherwise specified we used R version 4.3.2 (R Core Team 2019) for the rest of the analyses. For each locus in each island population, we then calculated the number of individuals typed, the number of alleles ( *A*, the raw count of alleles per locus), allelic richness (*A_r_*, following the extrapolation algorithm of Foulley and Ollivier (2006) as implemented in the *allelicrichness()* function from the R package pegas v. 1.3 (Paradis 2010)), observed and expected heterozygosity (*H_o_*, estimated as the number of heterozygous individuals compared to the total number of typed individuals, and *H_e_*, estimated using 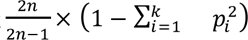 following Nei (1978)). We then tested if loci were in Hardy-Weinberg equilibrium (HWE) using a χ^2^ test and a permutation test based on 10,000 computer randomizations of genotypes resampled from the observed data. We summarized population-wise information over all loci using GenAlEx 6.5 (Peakall and Smouse 2012). Count of private alleles unbiased for sample size was calculated for each population using the formula 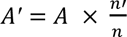, where *A* is the number of private alleles in a population, *n’* is the rarefied (minimum) sample size out of all populations, and *n* is the actual sample size of the population (Kalinowski 2004). We estimated rarefied allelic richness and pairwise F_ST_ between sampled islands using Weir and Cockerham (1984) as implemented in the *pairwise.WCfst()* function from the R package hierfstat v. 0.5–11 (Goudet 2005).

We investigated how abrupt reductions in population size may affect multi-locus heterozygosity (MLH) by calculating internal relatedness (IR; Amos et al. 2001). IR compares loci that are homozygous for rare alleles, assumed to be more likely associated with identity by descent, with loci that are homozygous for common alleles (Amos et al. 2001). We used IR instead of other indices to estimate MLH across the genome because IR operates well when average heterozygosity is low (Aparicio et al. 2006). We used the R package Rhh v. 1.0.2 (Alho et al. 2010) to calculate IR. Comparisons of IR across islands and age class (adults and hatchlings) were conducted using an Analysis of Variance (ANOVA) and a Tukey HSD test.

#### Bottleneck event signatures

Bottleneck detection is often difficult due to varying levels of pre-bottleneck genetic diversity, length and magnitude of the bottleneck, and immigration (Williamson-Natesan 2005). Nevertheless, populations that have recently undergone a bottleneck event present a specific distribution of allele frequencies such that only few alleles with frequencies < 0.1 are left in the population (Luikart and Cornuet 1998). Luikart and colleagues (1998) suggest that this particular signature can be detected using 5 to 20 loci and as little as 30 individuals. This signature can be detected for up to 80 generations after the bottleneck event (Cornuet and Luikart 1996; Luikart and Cornuet 1998). We used the software BOTTLENECK v1.2.02 (Cornuet and Luikart 1996; Piry et al. 1999) to estimate observed and expected heterozygosity excess across loci as a signature for genetic bottlenecks (Luikart and Cornuet 1998). We conducted 1000 iterations under the Infinite Allele Model (IAM) (Maruyama and Fuerst, 1985), the Stepwise Mutation Model (SMM) (Cornuet and Luikart 1996) and the Two-Phase Model (TPM) (Rienzo et al. 1994). TPM is considered an intermediate mutation model less extreme than IAM and SMM. We ran the TPM twice with separate parameter recommendations. The first was 90% of microsatellite mutations as stepwise, 10% as multi-step, an assuming a variance *_g_^2^* of 12 (Rienzo et al. 1994; Garza and Williamson 2001). The second was 78% of microsatellite mutations as stepwise, 22% as multi-step, and assuming a variance 𝜎*_g_^2^* of 12, which better represents vertebrate microsatellite mutation rates (Peery et al. 2012). Significant deviations from expected heterozygosity excess were determined with a Wilcoxon signed rank test and the L-shape distribution of allele frequency distribution (Cornuet and Luikart 1996; Piry et al. 1999). We also calculated the M-ratio (Garza and Williamson 2001) for each island using the R package strata G (Archer et al. 2017). The M-ratio uses the number of alleles and range of allele sizes at a locus to estimate the genetic bottleneck effect (Garza and Williamson 2001).

We performed an individual assignment test using the Bayesian algorithm implemented in STRUCTURE (Pritchard et al. 2000) to determine how the two serial bottleneck events have reshaped the distribution of genetic variation. We ran the analysis with a flat prior for the admixed parameter and we allowed the allele frequencies to be correlated among sampling sites. We tested values of K (number of genetic clusters) ranging from 1 to 4 and performed 20 replicates of the analysis for each value of K (Gilbert et al. 2012). STRUCTURE ran 10 ^6^ Markov Chain Monte Carlo (MCMC) in each analysis while discarding the first 10^5^ as burn-in. We used the command line back-end of STRUCTURE and the R package ParallelStructure v. 1.0 (Besnier and Glover 2013) to run the analysis. The output of this analysis is used to calculate the second order of differences in the likelihood function of K [ΔK] using the online tool Structure-Harvester (Evanno et al. 2005; Earl and VonHoldt 2012).

We further investigated the genetic structure of *C. pinguis* with a discriminant analysis of principal components (DAPC) (Jombart et al. 2010). As DAPC does not rely on any priors or model assumptions, unlike STRUCTURE, we can consider this approach as complementary to the Bayesian algorithm. We ran the analysis using the R package adegenet v. 2.1.10 (Jombart et al. 2008). We followed the author guidelines as implemented by others (Jombart et al. 2008, 2010; Welch et al. 2017; Pasachnik et al. 2020) to identify variables extrapolated from allele frequencies (principal components) and to cluster individuals, maximizing between group variation while minimizing variation within groups (Jombart et al. 2010).

Another consequence of population size fluctuation is the change in the degree of association between alleles at any two or more loci (*i.e.*, linkage disequilibrium or LD). In general, in small populations a higher correlation between alleles is expected, while little to no correlation is expected in large populations at equilibrium. We first estimated LD using the correlation coefficient, <u>𝑟</u>_d_, developed by Agapow and Burt (2001) and implemented in the *ia()* and *pair.ia()* functions from the R package poppr 2.9.6 (Kamvar et al. 2014; Kamvar et al. 2015). We then attempted to infer effective population size (*N_e_*) with NeEstimator V2.1 (Do et al. 2014). The software uses four different algorithms to infer *N_e_*: one based on linkage disequilibrium (Waples and Do 2008), one based on excess heterozygosity (Zhdanova and Pudovkin 2008), one based on molecular coancestry (Nomura 2008), and one based on the temporal variation in allele frequency, considering a lowest allele frequency of 0.02 (Nei and Tajima 1981; Pollak 1983; Jorde and Ryman 2007).

## Results

### Genetic diversity

We found no clear evidence of null alleles in our data set, nor consistent evidence of genotyping mistakes across loci and populations (van Oosterhout et al. 2004, Supplementary 2). While locus D137 showed an excess of homozygosity in the Anegada population, this was not true for the Guana or Necker populations. Loci D114, D130 and D135 showed excess homozygosity in individuals sampled from Guana Island while no loci showed any sign of genotyping errors from null alleles in Necker Island individuals (Supplementary Table 2). The algorithm used by FreeNA did not identify loci with a frequency of null alleles greater than 0.2 (max. value 0.118 for locus D137 in Anegada population, Supplementary Table 2), the threshold level considered for significance (Chapuis and Estoup 2007; Chapuis et al. 2008). Only 13 (ca. 19%) estimations were greater than 0.05, representing moderate null allele frequencies. D114 appears to have increased in null allele frequency by 10-fold after the first bottleneck event (Anegada: 0.010, Guana: 0.115, Necker: 0.097; Supplementary Table 2). There were no deviations from HWE both by loci and by population (Supplementary Table 3). We therefore decided to maintain all loci in the rest of the analyses.

Overall, the Anegada population showed greater diversity than Guana and Necker. The summary statistics calculated over all loci indicated a greater number of alleles as well as greater values of observed and expected heterozygosity in the source population as compared to the two sequentially translocated populations (Table 1). In particular, the level of genetic diversity (as expressed by expected heterozygosity) is 10.8% lower after the first translocation event, and an additional 10.3% is lost with the following translocation with an overall loss of diversity of 21% (Table 1).

We found increased population differentiation based on the variance in allele frequencies consistent with the sequential bottleneck events. Weir and Cockerham’s (1984) calculation of F_ST_ increased from 0.07 between Anegada and Guana to 0.13 between Anegada and Necker (Table 2).

**Table 2.** Population pairwise comparison of F_ST_ values as calculated following Weir and Cockerham (1984).

|  | Anegada | Guana | Necker |
| --- | --- | --- | --- |
| Anegada | NA |  |  |
| Guana | 0.068 | NA |  |
| Necker | 0.133 | 0.092 | NA |

Anegada internal relatedness values were significantly different from the Necker population (ANOVA, p - val = 0.034). IR increased sequentially after each bottleneck event, though not significantly (Figure 2). However, when looking only at the adults, there are no significant differences across populations (Figure 3A, ANOVA, p -val > 0.668). In hatchlings, there were significant differences between Anegada and Necker (Figure 3B, ANOVA, p -val = 0.013), but Guana did not significantly differ from either population. Hatchlings and adults from the same island did not significantly differ in IR from each other (Tukey HSD test, p-val > 0.10), though in general hatchlings had higher IR values than adults.

**Fig. 2.**
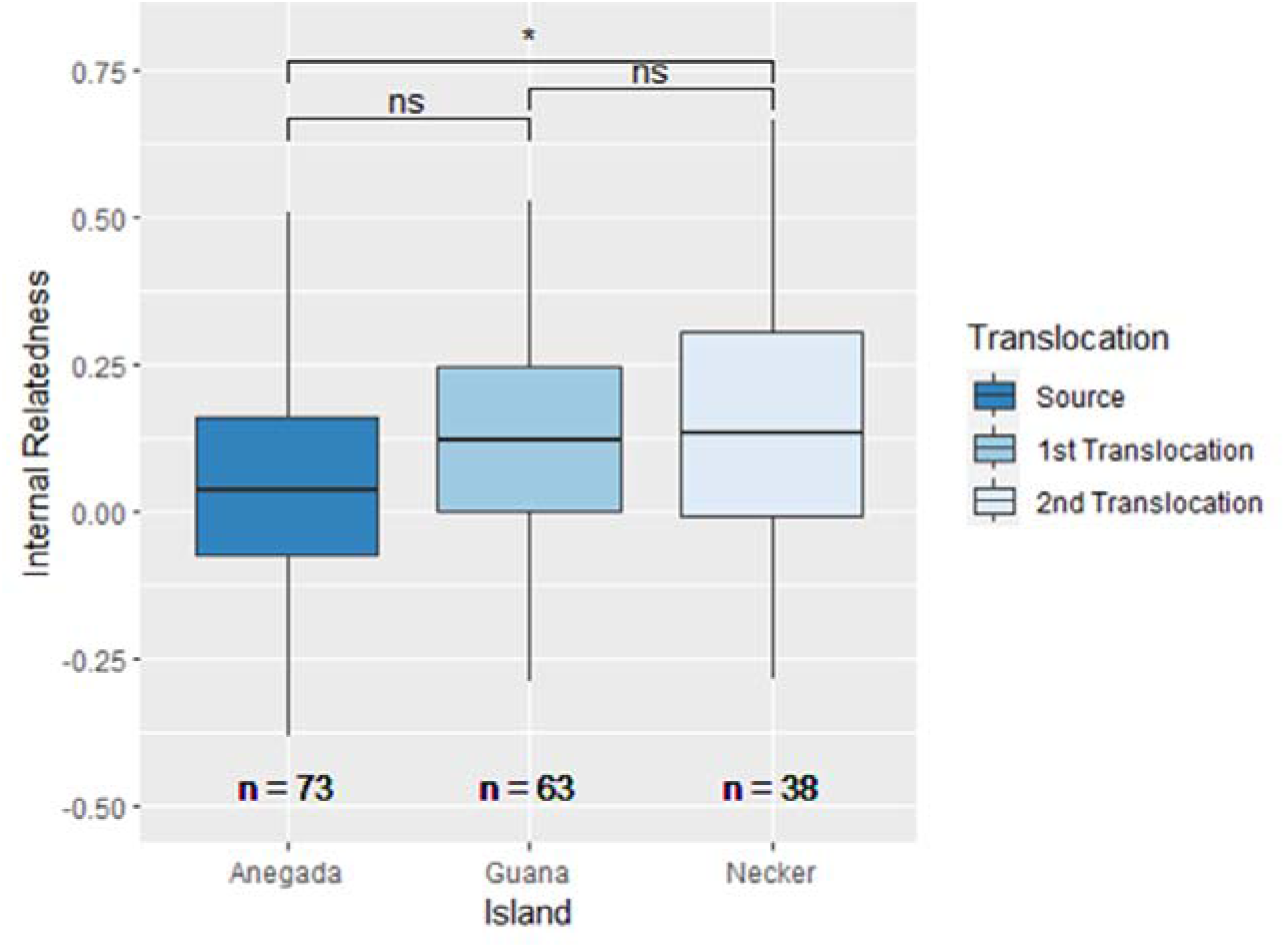
Internal relatedness values for Anegada, Guana, and Necker *Cyclura pinguis* populations. Sample sizes are below each box plot. * denotes p-values < 0.05 based on Tukey’s HSD, ns = not significant

**Fig. 3.**
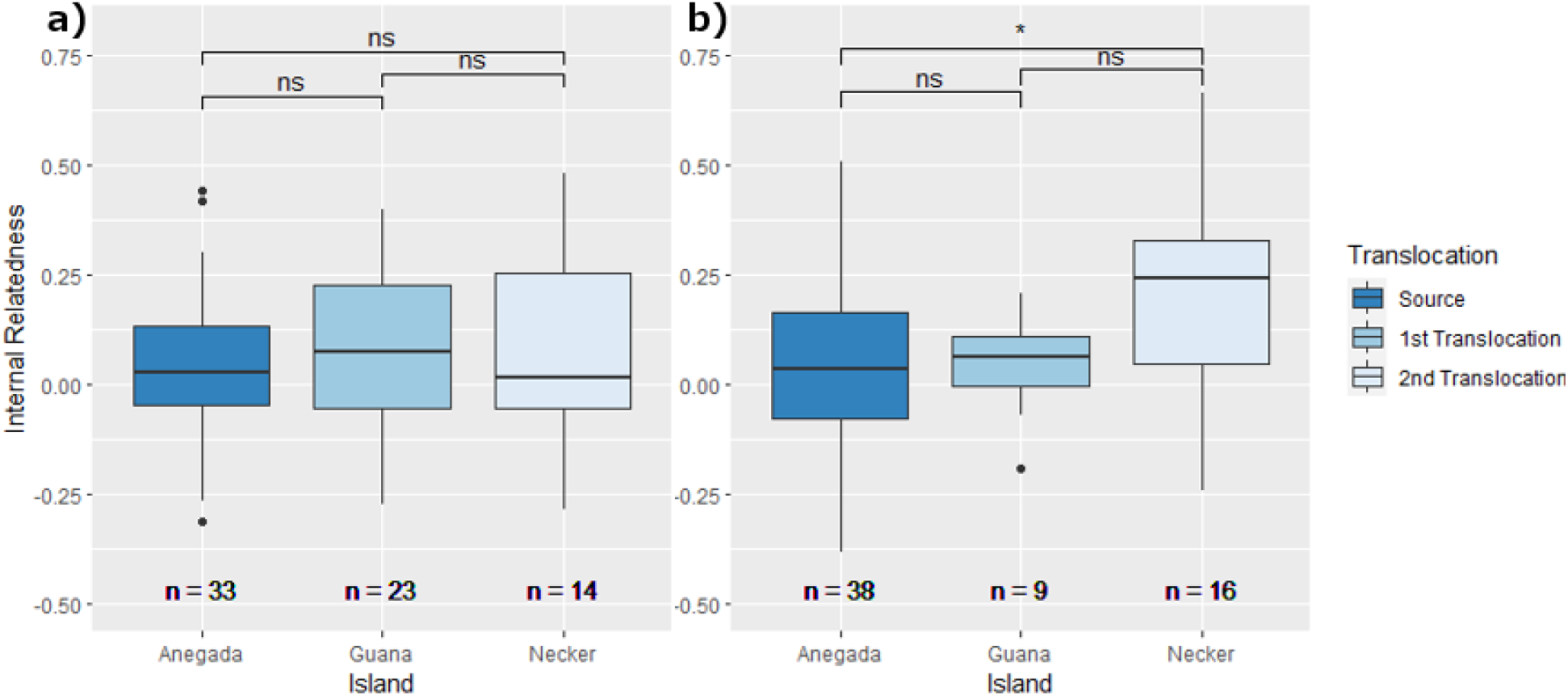
Internal relatedness values for Anegada, Guana, and Necker *Cyclura pinguis* populations split by age class, a) adults and b) hatchlings. Sample sizes are below each box plot. * denotes p < 0.05 based on Tukey’s HSD, ns = not significant

### Bottleneck event signatures

All tests under the Infinite Allele Model were statistically significant (p-val << 0.01) while none of the tests using the SMM were significant (Supplementary Table 4). Tests under the TPM were statistically significant (p-val < 0.05) for Guana but not for Necker, indicating only Guana had significant heterozygosity excess across loci after translocation events (Table 3A and 3B). Interestingly, following the parametrization proposed by Peery et al. (2012), Anegada also was found to have significant heterozygosity excess across loci (Table 3B). M -ratio on all three islands was in the range of populations that have not undergone a demographic bottleneck (M > 0.650) rather than those that have (M < 0.650) (Garza and Williamson 2001; Table 4).

**Table 3.** **a)** BOTTLENECK output from a Two-Phase Model (TPM) using a variance of 12, 10% step-wise mutations, and 1000 iterations based on Garza and Williamson 2001. **b)** BOTTLENECK output from a Two-Phase Model (TPM) using a variance of 12, 22% step-wise mutations, and 1000 iterations based on Peery et al. 2012. Significant values are bolded and in red.

|  | Island | Sign Test | 1-tail Wilcoxon | 2-tail Wilcoxon |
| --- | --- | --- | --- | --- |
| **a)** | Anegada | 0.519 | 0.108 | 0.216 |
|  | Guana | 0.051 | **0.013** | **0.026** |
|  | Necker | 0.335 | 0.226 | 0.452 |
| **b)** | Anegada | 0.456 | **0.015** | **0.029** |
|  | Guana | **0.044** | **0.003** | **0.007** |
|  | Necker | 0.091 | 0.084 | 0.168 |

**Table 4.** Values of Garza and Williamson (2001) M-ratio for Anegada, Guana, and Necker *Cylcura pinguis* populations. In general, populations that have undergone a genetic bottleneck have M < 0.650.

| Island | M-Ratio | Variance |
| --- | --- | --- |
| Anegada | 0.857 | 0.048 |
| Guana | 0.863 | 0.046 |
| Necker | 0.814 | 0.071 |

The second order of differences in the likelihood function of K [ΔK] indicated K = 3 as the most likely number of genetic clusters (Table 5). Individuals were grouped largely according to their sampling location (Figure 4). However, there was one individual from Guana whose genotype seemed to match the Necker cluster and two individuals from Necker whose genotypes matched more to the Guana cluster (Figure 4).

**Fig. 4.**
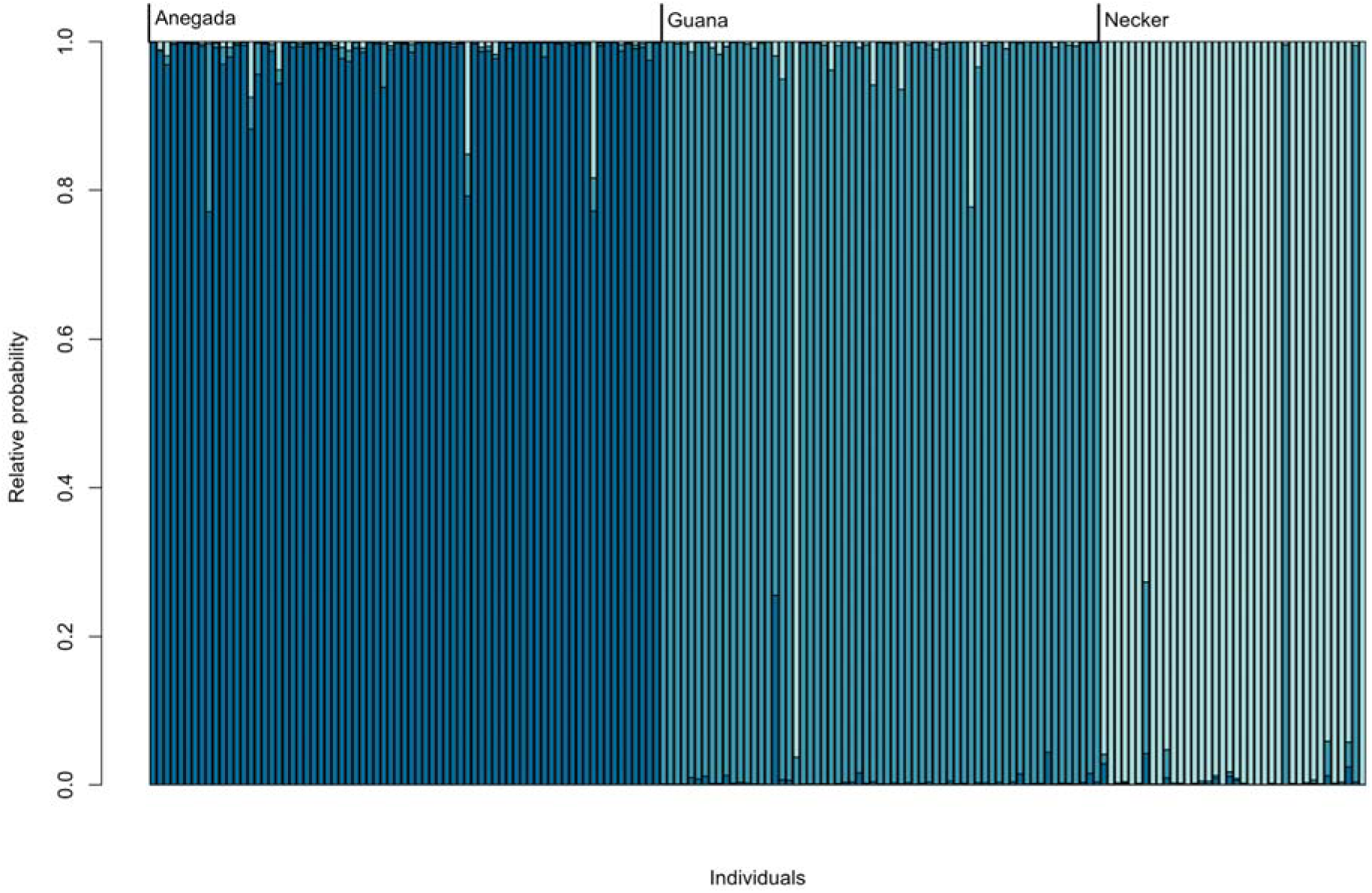
STRUCTURE results presented as a barplot. Individuals, represented as bars, are ordered by sampling site. The software groups them using colors based on the relative probability that, genetically, the individual belongs to a specific cluster

**Table 5.**
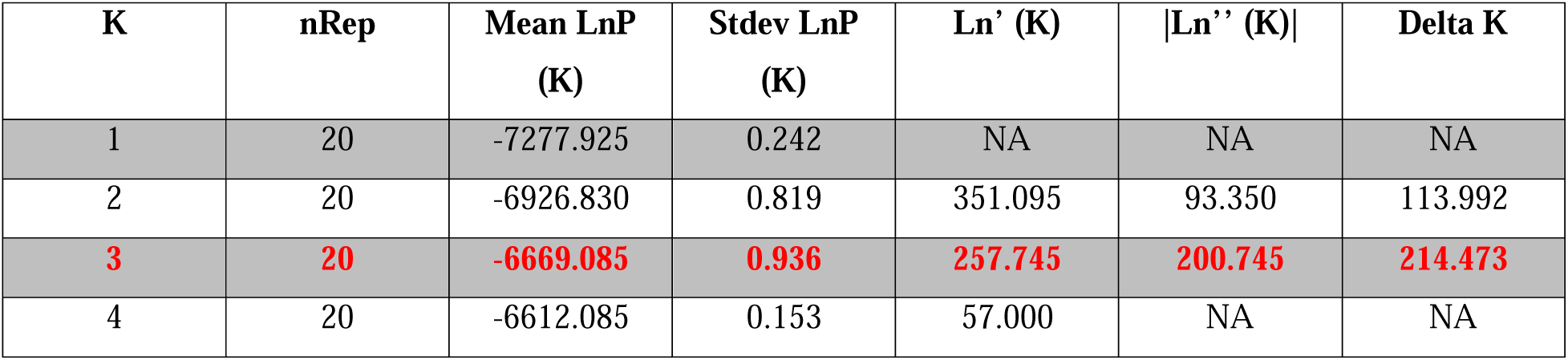
Table of the Evanno method output. Values highlighted in red show the largest value in the Delta K column, indicating that the number of K groups to best explain the data is 3.

A very similar result was obtained using the DAPC analysis. Figure 5 shows how individuals are clustered together along the two main principal components describing the differences in allele frequency between populations.

**Fig. 5.**
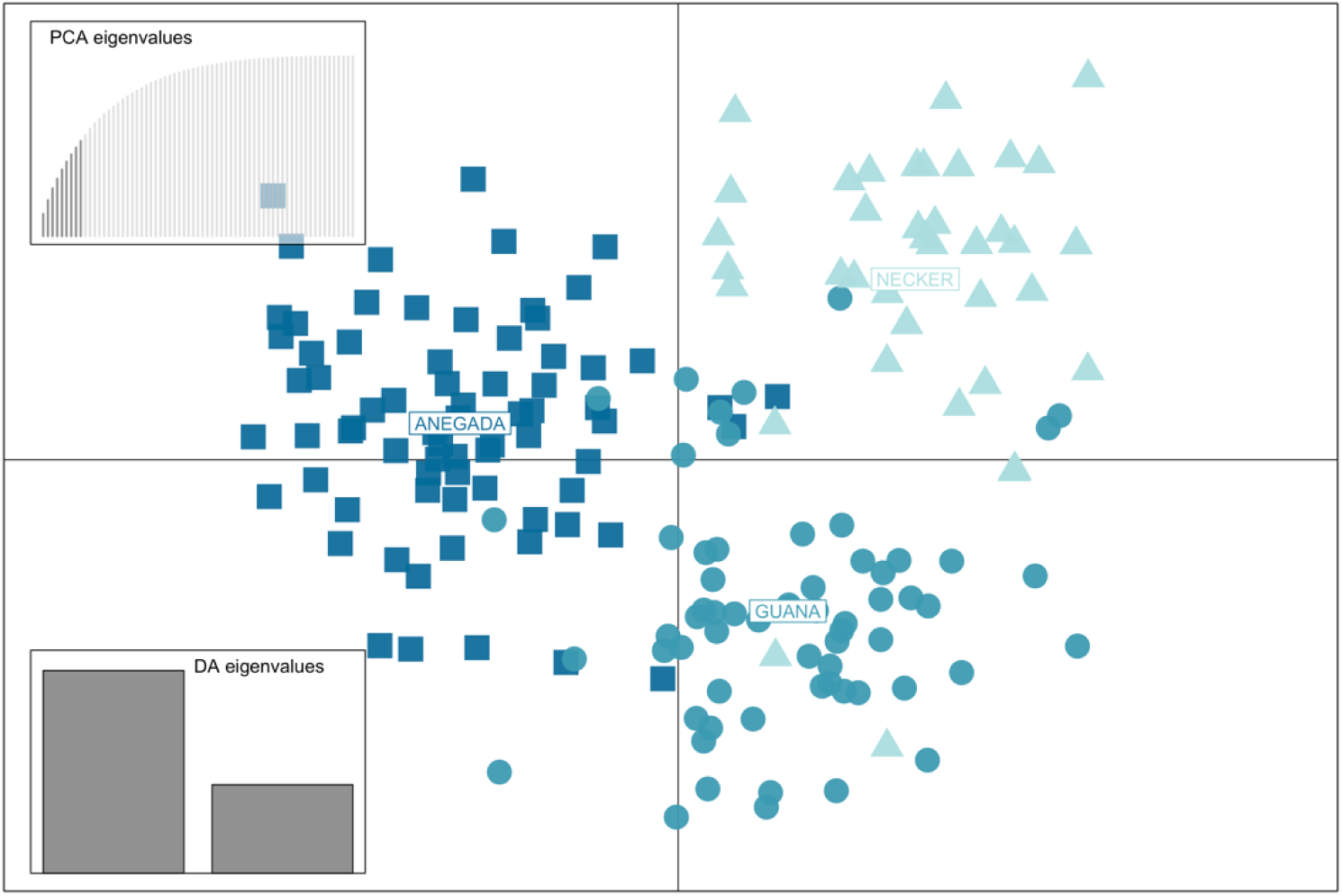
Discriminant Analysis of Principal Components. The scatterplot shows the first two principal components using sampling site as prior clusters, with samples from Anegada represented as squares, samples from Guana represented as circles, and samples from Necker represented as triangles. The two smaller plots display the Principal Component Analysis eigenvalues and the Discriminant Analysis eigenvalues, respectively

Estimates of linkage disequilibrium as expressed by the correlation coefficient <u>𝑟</u>_d_ were highest among individuals sampled on Necker Island (<u>𝑟</u>_d_ = 0.0360, p-val << 0.01). The second highest value was recorded from individuals sampled on Anegada (<u>𝑟</u>_d_ = 0.0113, p-val < 0.01) while Guana individuals showed the lowest, but still significant, indication of linkage disequilibrium (<u>𝑟</u>_d_ = 0.0111, p-val < 0.01; Table 6 and Supplementary Figures 1-3).

**Table 6.**
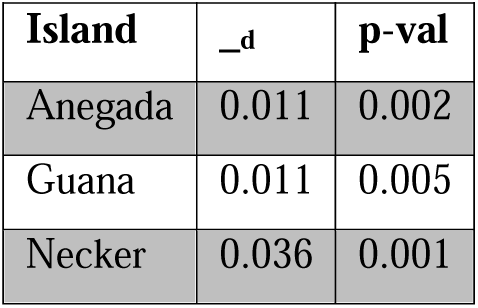
Analysis of linkage-disequilibrium. Sampling island, unbiased estimator of correlation among loci (<u>𝑟</u>_d_), results of significance test after 1000 permutations (p-val).

Our estimates of effective population size using the LD and heterozygote-excess methods are reported in Table 7. The results of these approaches are different from the demographic history we know through translocations. A better approximation of what we know has happened demographically is provided by the other two estimates. The molecular coancestry method estimated an infinite effective population size on Anegada, *N_e_* of 5.8 (3.4–8.9) on Guana, and 3.1 (1.3–5.7) on Necker. Finally, Nei and Tajima (1981)’s temporal method estimated two values of effective population size: *N_e_* = 6.9 (4.6–10.1) on Guana and *N_e_* = 5.1 (3.2–7.7) on Necker.

**Table 7.** Comparison of four methods (Linkage Disequilibrium –LD, Heterozygous Excess–Het. Exc., Coancestry and Temporal as implemented in the program NeEstimator) of calculating effective population size in *Cyclura pinguis*. Values of N_e_ are reported with the associated 95% confidence intervals.

| Island | LD | Het. Exc. | Coancestry | Temporal |
| --- | --- | --- | --- | --- |
| Anegada | 29.5 (25.2–34.8) | 34.6 (29.3–41.2) | Inf. | N/A |
| Guana | 36.8 (28.6–48.7) | 38.2 (29.6–50.9) | 5.8 (3.4–8.9) | 6.9 (4.6–10.1) |
| Necker | 25.6 (18.3–37.8) | 21.8 (16.3–30.3) | 3.1 (1.3–5.7) | 5.1 (3.2–7.7) |

## Discussion

In this manuscript we describe the genetic signature of two translocation events, with the second one seeded using individuals from the first. While we found evidence consistent with the occurrence of a genetic bottleneck and with the potential of an ongoing genetic paradox in both populations of *C. pinguis* established by translocation, we currently cannot ascertain the long-term adaptive repercussion of such genetic depletion.

Due to the small number of individuals moved during the two translocations, we anticipated that the genetic variation would be unevenly distributed across islands, with a progressive loss of variability as the translocations occurred. The first line of evidence supporting our prediction is provided by the documented decrease of average number of different alleles, observed, and expected heterozygosity from the source (Anegada), to the first (Guana) and the second (Necker) translocation. In particular, our results indicate that iguanas on Necker Island have experienced an overall loss of diversity, as measured by expected heterozygosity, greater than 20% when compared to the source population on Anegada (Table 1). This pattern is consistent with findings in other studies of translocations using small founding populations (Ewing et al. 2008; Jamieson 2011). The recommendation for maintaining an effective population size of 50 or more was established because theory indicates this should be sufficient to prevent a loss of genetic diversity of more than 1% in the first 100 generations (Franklin 1980; Frankham 1995). It is further projected that 20 founder genome equivalents should be sufficient for the maintenance of 90% of genetic variation (Soulé et al. 1986). Larger founding populations and supplementation have shown more success in maintaining genetic diversity (Jamieson 2011; White et al. 2018; White et al. 2020). However, each translocation is unique and species’ life history traits, current habitat characteristics, environmental stochasticity, and management goals should be modeled to ensure population success (White et al. 2020). The data presented here, together with the historical reconstruction of the translocations involving only a handful of individuals, provide a well corroborated explanation of the rapid loss of variability experienced throughout the *C. pinguis* translocation process.

Both individual assignment tests (STRUCTURE and DAPC, Fig. 4 and Fig. 5) indicate that the standing genetic variation of the translocated population has drifted enough from that of Anegada, that individuals can now be assigned to different islands based solely on their genotypes. Only three individuals were clustered in a different genetic cluster despite their geographic provenience (Figure 4). We can speculate that the two individuals collected on Necker Island and clustered as genetically belonging to the Guana group could have been two of the 4 original founders moved in 1995. Similarly, the one individual sampled on Guana and genetically clustered with individuals from Necker could retain alleles that are now predominant on Necker Island.

Further evidence corroborating our genetic diversity expectations is provided by the increase in IR values from 0.04 measured in samples from Anegada, to 0.147 measured in samples from Necker (Figure 2). Specifically, we found that hatchling internal relatedness increased significantly after translocation events while adult internal relatedness did not change (Figure 3). When samples were pooled, internal relatedness increased after translocation events, seemingly driven by the hatchlings. Though non-significant, hatchlings had higher internal relatedness values when compared to adults from the same island (data not shown). Increased levels of homozygosity in hatchlings relative to adults may indicate that hard selection (density-independent selection favoring advantageous traits in the current environment) or soft selection (density-dependent selection on a population’s relative fitness in a specific habitat) is acting on inbred individuals (Christiansen 1975; Whitlock 2002). An external factor in these environments may be preventing homozygous hatchlings from reaching adulthood, favoring outbreeding. Due to the relative recentness of both translocations, carrying capacity has not been reached and therefore intraspecific competition (soft selection) is unlikely to be yet occurring (Ho and Agrawal, 2012), suggesting that inbreeding depression may be driven by the expression of deleterious alleles as expressed in the mating between related individuals (hard selection). This is different from other Caribbean rock iguanas, which show more support for soft selection driving inbreeding depression (Berk 2013; Colosimo 2016; Moss et al. 2019). While this is an interesting and important result from a conservation perspective, we are aware that the difference between age classes in the detected IR values could also be explained by a lack of statistical power due to small sample sizes, and a more detailed analysis aimed at disentangling hard versus soft selection using a more appropriate sample size could be necessary.

Though demographic bottlenecks via translocation occurred twice in these *C. pinguis* populations, we were unable to pick up significant genetic bottleneck signatures using heterozygosity excess tests. Other studies have also generated non-significant results from well-documented, human-driven demographic bottlenecks (Aguilar et al. 2008; Henry et al. 2009), even in reptiles (Davy and Murphy, 2014; Bradke et al. 2021). The intermediate mutation model (TPM) showed significant heterozygosity excess across loci on Guana and in one instance Anegada, but not on Necker. Heterozygosity excess on Anegada may result from the ongoing demographic bottleneck that was initiated in the 20th century. Garza and Williamson’s M-ratio also did not indicate evidence of bottlenecks. Similar results were found with a translocated, insular population of elk (Hundertmark and Van Daele, 2010). The authors posit that the M-ratio, reliant on allele size, would lag behind genetic diversity loss and thus only pick up on bottleneck events multiple generations ago. Because our populations are 3-5 generations separated from translocation and sampling events, the genetic bottleneck may be too recent for the M-ratio to detect (Hundertmark and Van Daele, 2010).

Many factors, such as associative overdominance, influence distributions of alleles thereby impacting heterozygosity excess tests (Gilligan et al. 2005). Single-sample population genetic methods assume mutation-drift equilibrium (Peery et al. 2012). Due to the demographic history and small population sizes of these three iguana populations, it is highly unlikely that mutation is adding genetic variation as much as genetic drift is decreasing it (Whitlock 2000). Heterozygosity excess tests may lose statistical power if population size increases rapidly and panmixia increases (Hundertmark and Van Daele, 2010). Though this may in part explain our non-significant results for Necker, heterozygosity excess tests have been criticized for being sensitive to the starting parameters of the two - phase mutation model (Spong and Hellbord, 2002; Peery et al., 2012). We see this in our own results with only one of two sets of parameters showing heterozygosity excess on Anegada. Microsatellite mutation rates across vertebrates vary significantly but are difficult to measure for every study species (Peery et al. 2012). Researchers are forced to use parameter values recommended by the software’s authors or use averages that may be very different from actual rates (Peery et al. 2012). Bayesian approaches using coalescent theory and MCMC may be more accurate at indicating genetic signatures after a bottleneck (Peery et al. 2012).

In most cases, the estimates of effective population sizes generated by the various methodologies exceeded census numbers of the translocated populations at their founding (Table 7). The least accurate was the heterozygosity excess method estimating an infinite *N_e_* for all three islands. This technique has been shown to be imprecise due to the function’s need for >20 breeders, a large sample size, and its sensitivity to deviations from random mating (Luikart and Cornuet, 1999; Wang et al. 2016). The one-sample linkage disequilibrium method has been shown to detect changes in *N_e_* only 1-2 generations after abrupt population size changes, like bottlenecks, better than the two-sample temporal method (Antao et al. 2011, Waples 2024). Surprisingly, the LD method generated estimates inconsistent with the known history of these translocations. This may be due to the lack of linkage disequilibrium after the first translocation, which we also saw with the r_d_ values (Table 6). Internal relatedness and heterozygosity excess results suggest that there are more heterozygous individuals in our dataset than expected. This can decrease linkage disequilibrium estimates (Sabatti and Risch, 2002). Both the two-sample temporal method and the one-sample molecular coancestry method estimated effective population sizes similar to the founding population on the translocated islands. These calculations are less impacted by genetic drift and bottlenecks than other methods, and are theorized to be less affected by small, inbred populations (Nomura 2008). The temporal method is robust to non-random mating, population structure, age structure, and overlapping generations (Wang et al. 2016). We believe this serves as an anecdote highlighting the accuracy of the molecular coancestry method over other one-sample methods for estimating effective population size. However, it is recommended to factor in multiple, independent estimates of *N_e_* to get a more complete picture of the study population (Luikart and Cornuet, 1999).

Both translocated populations of *C. pinguis* studied here meet at least two of the criteria for a genetic paradox set by Estoup et al. (2016). First, we found ample evidence supporting a loss of genetic variability in the translocated populations compared to that in the source population. Second, estimates of census population size on both Guana and Necker are in the hundreds. This indicates rapid population growth and establishment following introductions of very few animals. However, it is more difficult to assess whether these populations are meeting the third criterion and adapting to novel environmental conditions in their introduced range. It is possible that adaptation has not yet happened but is actively occurring (Meek et al. 2023). It might also be argued that *C. pinguis* were preadapted to their translocated island habitat. Fossil evidence from Puerto Rico implies that the entire Puerto Rican Bank may have been home to *C. pinguis* or its ancestral species (MacLean 1982). Even so, there are no reports of *C. pinguis* naturally occurring on Guana or Necker. If these *C. pinguis* populations successfully navigate potential pitfalls, they serve as a good example of a genetic paradox, and this may bode well for the future of translocation as a conservation strategy.

We conclude that the iguana populations of Guana and Necker are genetically depauperate and should not be considered as source populations for future conservation translocations. Though currently impacted by anthropogenic disturbances, the naturally occurring population of *C. pinguis* on Anegada remains the most appropriate genetic stock for future translocations. However, supplementing the populations on Guana and Necker with iguanas from Anegada may instigate genetic rescue (eg. Pimm et al. 2006). The decrease in genetic diversity on Guana and Necker is directly tied to the small number of founders translocated to each island (N=8 and N=4, respectively). The specific number of individuals translocated for conservation purposes depends heavily on the species, habitat, and other external factors (Furlan et al. 2020). However, larger founding populations decrease the impacts of genetic drift and prevent inbreeding depression, thereby preserving genetic diversity over multiple generations (Frankham et al. 2014). Though considered successfully colonized from a demographic perspective, we posit the populations of Guana and Necker represent a genetic paradox, though it is too early to comment on population fitness and evolution potential. Preventing inbreeding depression should be a high priority for wildlife managers considering conservation translocations to address long-term goals, such as population sustainability without human intervention. Conservation translocations can protect genetic diversity, if done thoughtfully, to ensure long-term resiliency through evolutionary time, especially in the face of a stochastic and rapidly changing world.

## Data Availability

R script and microsatellite genotypes associated with this project are available upon request.

## Compliance with Ethical Standards

All animal capture, handling, and sampling was performed following the American Society of Ichthyologists and Herpetologists (ASIH) guidelines for use of reptiles and amphibians in research and all methods were approved under the authors’ IACUC permits.

## Author Contributions

GC, MW and GPG designed the research; GP, ZTH, and GPG collected the samples; ZTH performed lab work; GC and ZD performed data analysis with guidance from MW, GG, and GPG; GC and ZD wrote the primary manuscript; all authors edited and revised the manuscript.

## Acknowledgments

We thank the owners of Guana Island and Necker Island for access to their islands and their staff and James Lazell for their assistance. This project was partially funded by The Conservation Agency through a grant from the Falconwood Foundation. Funding and laboratory space for this project was provided by San Diego Zoo Wildlife Alliance, with special thanks to Heidi Davis and William Modi. Our appreciation goes out to Nancy Pascoe (National Parks Trust of the Virgin Islands), Kelly Bradley (Fort Worth Zoo), and for the headstart program facility staff for sample collection and ongoing support of iguana conservation efforts.

