## Supplementary Information for "The impact of serial translocations on the genetic diversity of Anegada iguanas (*Cyclura pinguis*) in the British Virgin Islands"

**Supplementary Table 1:** List of 21 microsatellite loci used in this study. A=number of alleles, missing=number of missing genotypes, N=total number of samples, Obs. H= observed heterozygosity, Exp. H=expected heterozygosity.

| **Name** | **Source** | **A** | **Missing** | **N** | **Obs. H** | **Exp. H** |
| --- | --- | --- | --- | --- | --- | --- |
| C6 | Lau et al. 2009 | 3 | 0 | 174 | 0.483 | 0.569 |
| C103 | Lau et al. 2009 | 5 | 0 | 174 | 0.575 | 0.64 |
| D11 | Lau et al. 2009 | 4 | 0 | 174 | 0.356 | 0.405 |
| D101 | Lau et al. 2009 | 4 | 0 | 174 | 0.466 | 0.518 |
| D102 | Lau et al. 2009 | 6 | 0 | 174 | 0.414 | 0.458 |
| D107 | Lau et al. 2009 | 4 | 0 | 174 | 0.638 | 0.672 |
| D110 | Lau et al. 2009 | 4 | 0 | 174 | 0.575 | 0.627 |
| D114 | Lau et al. 2009 | 4 | 0 | 174 | 0.414 | 0.527 |
| D120 | Lau et al. 2009 | 6 | 0 | 174 | 0.707 | 0.706 |
| D124 | Lau et al. 2009 | 3 | 0 | 174 | 0.621 | 0.62 |
| D130 | Lau et al. 2009 | 3 | 0 | 174 | 0.402 | 0.53 |
| D135 | Lau et al. 2009 | 3 | 0 | 174 | 0.454 | 0.58 |
| D136 | Lau et al. 2009 | 6 | 0 | 174 | 0.684 | 0.672 |
| D140 | Lau et al. 2009 | 3 | 0 | 174 | 0.54 | 0.598 |
| C113 | Lau et al. 2009 | 3 | 0 | 174 | 0.201 | 0.227 |
| C124 | Lau et al. 2009 | 5 | 0 | 174 | 0.718 | 0.731 |
| D1 | Lau et al. 2009 | 3 | 0 | 174 | 0.213 | 0.261 |
| D9 | Lau et al. 2009 | 5 | 0 | 174 | 0.603 | 0.585 |
| D105 | Lau et al. 2009 | 4 | 0 | 174 | 0.621 | 0.667 |
| D111 | Lau et al. 2009 | 7 | 0 | 174 | 0.649 | 0.666 |
| D137 | Lau et al. 2009 | 4 | 0 | 174 | 0.264 | 0.366 |

**Supplementary Table 2:** Null allele estimates of all population and locus combinations from FreeNa (Chapuis and Estoup 2007) and Micro-Checker 2.2.3 (van Oosterhout et al. 2004), in order of population and highest null allele frequency estimate based on Chapuis and Estoup. Null alleles were determined present by Micro-Checker using van Oosterhout et al. 2004, Chakraborty et al. 1992, and Brookfield 1996 and highlighted in yellow.

| **Locus** | **Pop** | **Null Present** | **Chapuis & Estoup** | **Van Oosterhout** | **Chakraborty** | **Brookfield 1** | **Brookfield 2** |
| --- | --- | --- | --- | --- | --- | --- | --- |
| D137 | Anegada | yes | 0.1176 | 0.1500 | 0.1905 | 0.1072 | 0.1072 |
| C113 | Anegada | no | 0.0488 | 0.0618 | 0.0735 | 0.0330 | 0.0330 |
| C103 | Anegada | no | 0.0479 | 0.0660 | 0.0698 | 0.0512 | 0.0512 |
| D135 | Anegada | no | 0.0377 | 0.0264 | 0.0343 | 0.0242 | 0.0242 |
| D1 | Anegada | no | 0.0373 | 0.0627 | 0.0704 | 0.0361 | 0.0361 |
| D130 | Anegada | no | 0.0334 | 0.0081 | 0.0141 | 0.0088 | 0.0088 |
| D110 | Anegada | no | 0.0229 | 0.0350 | 0.0363 | 0.0279 | 0.0279 |
| D11 | Anegada | no | 0.0172 | 0.0307 | 0.0269 | 0.0103 | 0.0103 |
| D107 | Anegada | no | 0.0146 | 0.0287 | 0.0255 | 0.0206 | 0.0206 |
| D124 | Anegada | no | 0.0132 | 0.0039 | 0.0082 | 0.0065 | 0.0065 |
| D114 | Anegada | no | 0.0105 | 0.0087 | 0.0110 | 0.0081 | 0.0081 |
| D120 | Anegada | no | 0.0088 | 0.0452 | 0.0377 | 0.0297 | 0.0297 |
| D136 | Anegada | no | 0.0001 | -0.0036 | -0.0005 | -0.0004 | 0.0000 |
| D140 | Anegada | no | 0.0000 | -0.0183 | -0.0162 | -0.0128 | 0.0000 |
| C6 | Anegada | no | 0.0000 | -0.0244 | -0.0241 | -0.0118 | 0.0000 |
| D102 | Anegada | no | 0.0000 | 0.0001 | 0.0035 | 0.0026 | 0.0026 |
| C124 | Anegada | no | 0.0000 | -0.0233 | -0.0204 | -0.0179 | 0.0000 |
| D101 | Anegada | no | 0.0000 | -0.0823 | -0.0698 | -0.0508 | 0.0000 |
| D9 | Anegada | no | 0.0000 | -0.0814 | -0.0595 | -0.0466 | 0.0000 |
| D105 | Anegada | no | 0.0000 | -0.0188 | -0.0243 | -0.0197 | 0.0000 |
| D111 | Anegada | no | 0.0000 | -0.0141 | -0.0200 | -0.0166 | 0.0000 |
| D114 | Guana | yes | 0.1153 | 0.1657 | 0.2141 | 0.1199 | 0.1199 |
| D135 | Guana | yes | 0.1057 | 0.1481 | 0.1804 | 0.1023 | 0.1023 |
| D130 | Guana | yes | 0.0871 | 0.1301 | 0.1575 | 0.0935 | 0.0935 |
| D101 | Guana | no | 0.0663 | 0.1009 | 0.1164 | 0.0678 | 0.0678 |
| D137 | Guana | no | 0.0633 | 0.0858 | 0.1798 | 0.0255 | 0.0255 |
| D11 | Guana | no | 0.0500 | 0.0702 | 0.0868 | 0.0388 | 0.0388 |
| D1 | Guana | no | 0.0360 | 0.0518 | 0.0701 | 0.0167 | 0.0167 |
| D9 | Guana | no | 0.0299 | 0.0225 | 0.0308 | 0.0226 | 0.0226 |
| D140 | Guana | no | 0.0238 | 0.0351 | 0.0371 | 0.0232 | 0.0232 |
| D107 | Guana | no | 0.0198 | 0.0521 | 0.0488 | 0.0365 | 0.0365 |
| C113 | Guana | no | 0.0003 | -0.0011 | -0.0011 | -0.0004 | 0.0000 |
| C103 | Guana | no | 0.0002 | -0.0648 | -0.0493 | -0.0373 | 0.0000 |
| D102 | Guana | no | 0.0000 | 0.0059 | 0.0056 | 0.0035 | 0.0035 |
| C6 | Guana | no | 0.0000 | -0.0053 | -0.0127 | -0.0100 | 0.0000 |
| D110 | Guana | no | 0.0000 | -0.0160 | -0.0211 | -0.0147 | 0.0000 |
| D105 | Guana | no | 0.0000 | -0.0391 | -0.0327 | -0.0264 | 0.0000 |
| D111 | Guana | no | 0.0000 | 0.0150 | 0.0070 | 0.0049 | 0.0049 |
| D120 | Guana | no | 0.0000 | -0.0515 | -0.0438 | -0.0386 | 0.0000 |
| D124 | Guana | no | 0.0000 | -0.0589 | -0.0522 | -0.0413 | 0.0000 |
| D136 | Guana | no | 0.0000 | -0.0871 | -0.0650 | -0.0529 | 0.0000 |
| C124 | Guana | no | 0.0000 | -0.0267 | -0.0258 | -0.0225 | 0.0000 |
| D114 | Necker | no | 0.0971 | 0.1355 | 0.1891 | 0.0820 | 0.0820 |
| D1 | Necker | no | 0.0774 | 0.0809 | 0.1294 | 0.0334 | 0.0334 |
| C103 | Necker | no | 0.0619 | 0.0613 | 0.0678 | 0.0477 | 0.0477 |
| C6 | Necker | no | 0.0501 | 0.0255 | 0.0305 | 0.0206 | 0.0206 |
| D101 | Necker | no | 0.0483 | 0.0698 | 0.0796 | 0.0445 | 0.0445 |
| D130 | Necker | no | 0.0286 | 0.0419 | 0.0459 | 0.0239 | 0.0239 |
| D140 | Necker | no | 0.0140 | 0.0208 | 0.0215 | 0.0139 | 0.0139 |
| D105 | Necker | no | 0.0131 | 0.0412 | 0.0372 | 0.0214 | 0.0214 |
| D102 | Necker | no | 0.0001 | -0.0132 | -0.0066 | -0.0003 | 0.0000 |
| C113 | Necker | no | 0.0001 | -0.0132 | -0.0066 | -0.0003 | 0.0000 |
| D137 | Necker | no | 0.0000 | -0.0418 | -0.0185 | -0.0114 | 0.0000 |
| D110 | Necker | no | 0.0000 | -0.0489 | -0.0435 | -0.0306 | 0.0000 |
| D120 | Necker | no | 0.0000 | -0.0552 | -0.0444 | -0.0358 | 0.0000 |
| D135 | Necker | no | 0.0000 | 0.0077 | -0.0107 | -0.0069 | 0.0000 |
| C124 | Necker | no | 0.0000 | 0.0521 | 0.0325 | 0.0219 | 0.0219 |
| D11 | Necker | no | 0.0000 | -0.0853 | -0.0745 | -0.0625 | 0.0000 |
| D107 | Necker | no | 0.0000 | -0.0372 | -0.0265 | -0.0209 | 0.0000 |
| D124 | Necker | no | 0.0000 | -0.0090 | -0.0377 | -0.0248 | 0.0000 |
| D136 | Necker | no | 0.0000 | -0.0711 | -0.0600 | -0.0470 | 0.0000 |
| D9 | Necker | no | 0.0000 | -0.1664 | -0.1041 | -0.0720 | 0.0000 |
| D111 | Necker | no | 0.0000 | -0.1193 | -0.0900 | -0.0737 | 0.0000 |

**Supplementary Table 3:** Testing for Hardy-Weinberg Equilibrium using a χ^2^ test and a permutation test based on 10,000 computer randomizations of genotypes resampled from the observed data., separated by population. No loci in each population had a p-value < 0.05.

| **Loci** | **Pop** | **A** | **Obs. H** | **Exp. H** | **Chi sq** | **P-val** | **F** |
| --- | --- | --- | --- | --- | --- | --- | --- |
| C6 | Anegada | 3 | 0.329 | 0.313 | 14.012 | 0.999 | -0.051 |
| C103 | Anegada | 4 | 0.562 | 0.646 | 31.388 | 1.000 | 0.130 |
| D11 | Anegada | 3 | 0.233 | 0.246 | 10.146 | 0.994 | 0.053 |
| D101 | Anegada | 4 | 0.589 | 0.512 | 44.998 | 1.000 | -0.150 |
| D102 | Anegada | 6 | 0.589 | 0.593 | 36.972 | 1.000 | 0.007 |
| D107 | Anegada | 4 | 0.671 | 0.706 | 34.708 | 1.000 | 0.050 |
| D110 | Anegada | 4 | 0.616 | 0.663 | 44.558 | 1.000 | 0.071 |
| D114 | Anegada | 4 | 0.575 | 0.588 | 23.381 | 1.000 | 0.022 |
| D120 | Anegada | 6 | 0.644 | 0.694 | 38.67 | 1.000 | 0.072 |
| D124 | Anegada | 3 | 0.644 | 0.655 | 35.166 | 1.000 | 0.017 |
| D130 | Anegada | 3 | 0.452 | 0.465 | 22.499 | 1.000 | 0.028 |
| D135 | Anegada | 3 | 0.534 | 0.572 | 19.703 | 1.000 | 0.066 |
| D136 | Anegada | 5 | 0.685 | 0.684 | 36.623 | 1.000 | -0.001 |
| D140 | Anegada | 3 | 0.658 | 0.637 | 26.746 | 1.000 | -0.033 |
| C113 | Anegada | 3 | 0.274 | 0.317 | 8.452 | 0.962 | 0.136 |
| C124 | Anegada | 5 | 0.781 | 0.75 | 43.493 | 1.000 | -0.041 |
| D1 | Anegada | 3 | 0.329 | 0.379 | 9.109 | 0.989 | 0.132 |
| D9 | Anegada | 5 | 0.658 | 0.584 | 62.181 | 1.000 | -0.127 |
| D105 | Anegada | 4 | 0.685 | 0.652 | 39.998 | 1.000 | -0.051 |
| D111 | Anegada | 7 | 0.712 | 0.684 | 62.535 | 1.000 | -0.041 |
| D137 | Anegada | 4 | 0.342 | 0.504 | 23.427 | 1.000 | 0.321 |
| C6 | Guana | 3 | 0.651 | 0.634 | 28.503 | 1.000 | -0.027 |
| C103 | Guana | 4 | 0.619 | 0.561 | 31.819 | 1.000 | -0.103 |
| D11 | Guana | 3 | 0.27 | 0.321 | 6.472 | 0.989 | 0.159 |
| D101 | Guana | 3 | 0.381 | 0.481 | 9.581 | 0.998 | 0.208 |
| D102 | Guana | 4 | 0.444 | 0.449 | 21.01 | 1.000 | 0.011 |
| D107 | Guana | 4 | 0.587 | 0.648 | 20.572 | 0.999 | 0.094 |
| D110 | Guana | 4 | 0.54 | 0.517 | 24.055 | 1.000 | -0.044 |
| D114 | Guana | 3 | 0.333 | 0.515 | 8.745 | 0.987 | 0.353 |
| D120 | Guana | 5 | 0.794 | 0.727 | 42.886 | 1.000 | -0.092 |
| D124 | Guana | 3 | 0.667 | 0.601 | 30.603 | 1.000 | -0.110 |
| D130 | Guana | 3 | 0.381 | 0.523 | 9.879 | 0.993 | 0.272 |
| D135 | Guana | 3 | 0.349 | 0.503 | 12.015 | 0.993 | 0.306 |
| D136 | Guana | 4 | 0.698 | 0.613 | 44.439 | 1.000 | -0.139 |
| D140 | Guana | 2 | 0.444 | 0.479 | 16.371 | 1.000 | 0.073 |
| C113 | Guana | 2 | 0.222 | 0.222 | 7.048 | 0.992 | 0.000 |
| C124 | Guana | 4 | 0.778 | 0.739 | 36.607 | 1.000 | -0.053 |
| D1 | Guana | 2 | 0.127 | 0.146 | 3.43 | 0.936 | 0.130 |
| D9 | Guana | 3 | 0.571 | 0.608 | 18.254 | 1.000 | 0.061 |
| D105 | Guana | 3 | 0.683 | 0.639 | 33.83 | 1.000 | -0.069 |
| D111 | Guana | 4 | 0.54 | 0.547 | 24.49 | 1.000 | 0.013 |
| D137 | Guana | 3 | 0.063 | 0.091 | 8.929 | 0.997 | 0.308 |
| C6 | Necker | 3 | 0.5 | 0.532 | 19.149 | 1.000 | 0.060 |
| C103 | Necker | 4 | 0.526 | 0.603 | 15.144 | 0.981 | 0.128 |
| D11 | Necker | 3 | 0.737 | 0.635 | 22.596 | 1.000 | -0.161 |
| D101 | Necker | 2 | 0.368 | 0.432 | 4.551 | 0.967 | 0.148 |
| D102 | Necker | 2 | 0.026 | 0.026 | 0.52 | 1.000 | 0.000 |
| D107 | Necker | 4 | 0.658 | 0.624 | 25.629 | 1.000 | -0.054 |
| D110 | Necker | 4 | 0.553 | 0.507 | 19.038 | 1.000 | -0.091 |
| D114 | Necker | 2 | 0.237 | 0.347 | 3.383 | 0.934 | 0.317 |
| D120 | Necker | 4 | 0.684 | 0.626 | 24.775 | 1.000 | -0.093 |
| D124 | Necker | 3 | 0.5 | 0.464 | 19.961 | 1.000 | -0.078 |
| D130 | Necker | 2 | 0.342 | 0.375 | 5.027 | 0.975 | 0.088 |
| D135 | Necker | 3 | 0.474 | 0.464 | 12.461 | 0.998 | -0.022 |
| D136 | Necker | 5 | 0.658 | 0.583 | 42.257 | 1.000 | -0.129 |
| D140 | Necker | 2 | 0.474 | 0.494 | 13.692 | 0.999 | 0.040 |
| C113 | Necker | 2 | 0.026 | 0.026 | 0.52 | 1.000 | 0.000 |
| C124 | Necker | 5 | 0.5 | 0.534 | 20.907 | 1.000 | 0.064 |
| D1 | Necker | 3 | 0.132 | 0.171 | 39.325 | 1.000 | 0.228 |
| D9 | Necker | 3 | 0.553 | 0.448 | 22.243 | 1.000 | -0.234 |
| D105 | Necker | 3 | 0.395 | 0.425 | 7.158 | 0.972 | 0.071 |
| D111 | Necker | 4 | 0.711 | 0.593 | 26.83 | 1.000 | -0.199 |
| D137 | Necker | 3 | 0.447 | 0.431 | 12.697 | 1.000 | -0.037 |

**Supplementary Table 4:** Results of BOTTLENECK software using the Infinite Allele Model (IAM) and the Stepwise Mutation Model (SMM) with default parameters. None of the SMM tests were statistically significant while all IAM tests were statistically significant.

|  | **Island** | **Sign Test** | **1-tail Wilcoxon** | **2-tail Wilcoxon** |
| --- | --- | --- | --- | --- |
| **IAM** | Anegada | 0.00004 | 0.000001 | 0.000002 |
|  | Guana | 0.00012 | 0.000033 | 0.000065 |
|  | Necker | 0.00072 | 0.002439 | 0.004879 |
| **SMM** | Anegada | 0.33677 | 0.50000 | 1.00000 |
|  | Guana | 0.26472 | 0.06864 | 0.13728 |
|  | Necker | 0.53669 | 0.52713 | 0.97286 |

**Supplementary Figure 1:** Heat map of correlation coefficient values $\underline{r}$_d_ (index of association) between all pairwise combinations of 21 microsatellite loci for the Anegada population.

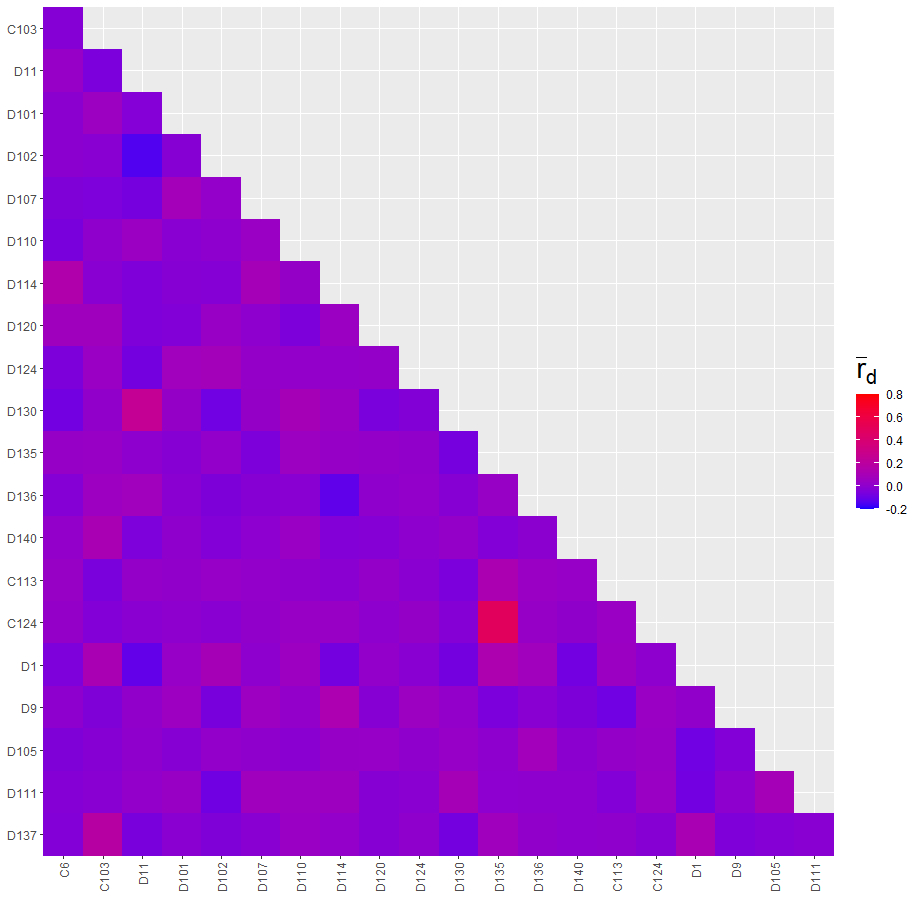

**Supplementary Figure 2:** Heat map of correlation coefficient values $\underline{r}$_d_ (index of association) between all pairwise combinations of 21 microsatellite loci for the Guana population.

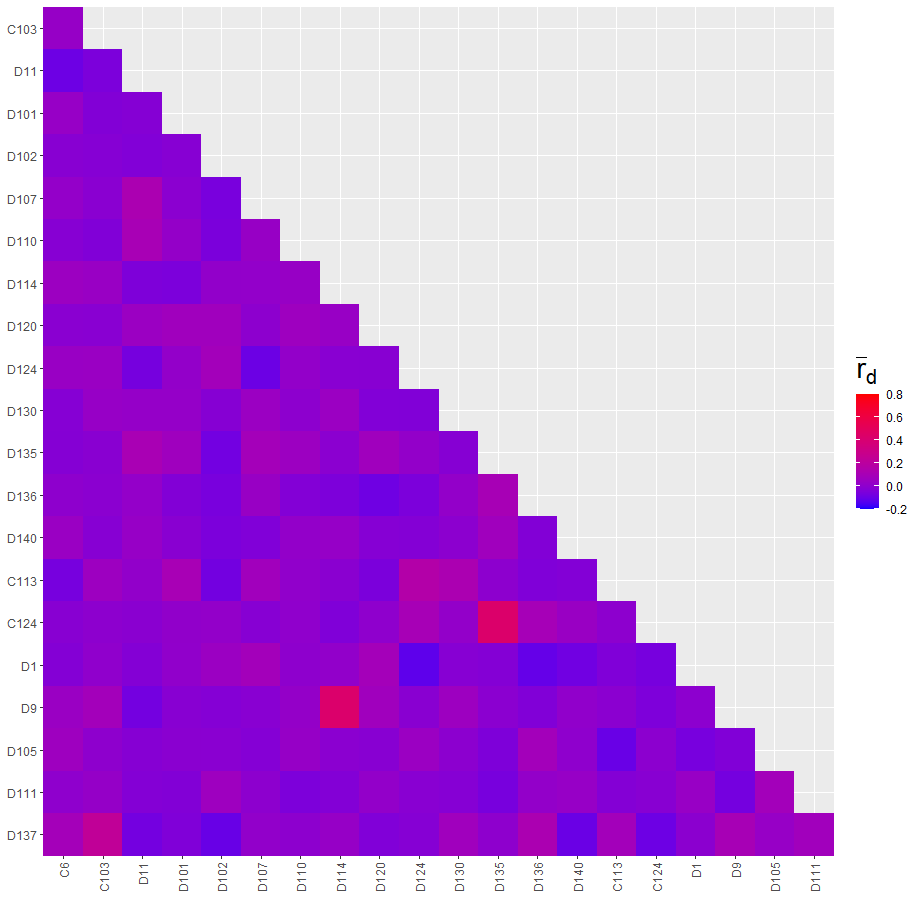

**Supplementary Figure 3:** Heat map of correlation coefficient values $\underline{r}$_d_ (index of association) between all pairwise combinations of 21 microsatellite loci for the Necker population.

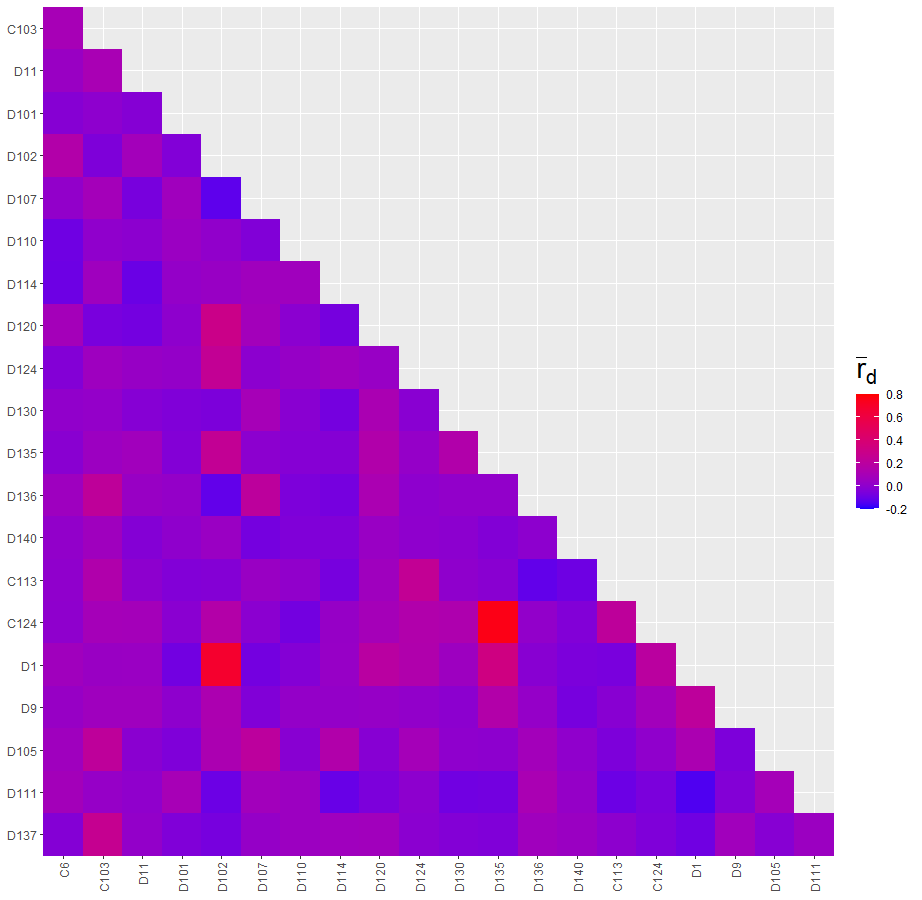
